# Challenging the Conventional Treatment Initiation Paradigm: Early Detection of Irreversible Cellular Damage in Cardiac Biopsies of Fabry Disease Before the Formation of Gb3 Inclusion Bodies

**DOI:** 10.1101/2024.06.28.601296

**Authors:** Chung-Lin Lee, Pei-Sin Chen, Yu-Ying Lu, Yu-Ting Chiang, Ching-Tzu Yen, Chun-Ying Huang, Yen-Fu Cheng, Hsiang-Yu Lin, Yun-Ru Chen, Dau-Ming Niu

**Author notes:** Prof. Dau-Ming Niu, Department of Pediatrics, Taipei Veterans General Hospital, No.201, Sec. 2, Shipai Rd., Beitou District, Taipei City, Taiwan 11217, R.O.C., Dr. Yun-Ru Chen, Department of Pediatrics, Taipei Veterans General Hospital, No.201, Sec. 2, Shipai Rd., Beitou District, Taipei City, Taiwan 11217, R.O.C.

## Abstract

**Background:** Fabry disease (FD) is a lysosomal storage disorder impacting multiple organs, including the heart. We investigated whether early-stage globotriaosylceramide (Gb3) accumulation, before occurrence of inclusion bodies, could cause significant stress and irreversible damages of the cardiomyocytes in FD patients. To assess the cellular stress and irreversible damage of cardiomyocytes in FD during early-stage Gb3 accumulation before the occurrence of typical pathology.

**Methods:** Immunofluorescent (IF) staining or Western blotting were performed on fibroblasts from FD patients and myocardial biopsies from G3Stg/GLAko mice and FD patients. Notably, all biopsies exhibited detectable Gb3 accumulation under IF but lacked typical FD (Gb3 inclusion body) pathology. Staining targeted nuclear factor-κB (NF-κB), interleukin-18 (IL-18), phospho-p42/44 mitogen-activated protein kinase (MAPK), and inducible nitric oxide synthase (iNOS) as inflammatory and oxidative stress markers. Alpha-smooth muscle actin (α-SMA) IF staining was conducted to detect myofibroblasts.

**Results:** Fibroblasts from FD patients, in conjunction with cardiomyocytes from both G3Stg/GLAko mice and FD patients, exhibited significant accumulation of inflammatory markers such as NF-κB IL-18 and phospho-p42/44 MAPK, as well as the oxidative stress marker iNOS. Despite the absence of typical FD pathology, the presence of fibrosis was confirmed in myocardial biopsies from these patients through strong positive staining of α-SMA.

**Conclusions:** Significant cellular stress and even irreversible damage may occur before the onset of typical pathological changes in cardiomyocytes of FD. Based on our findings, treatment should be initiated much earlier than we currently thought to prevent irreversible damage and improve the prognosis of FD patients.

## Introduction

Fabry disease (MIM 301500) is an X-linked lysosomal storage disorder due to deficient alpha-galactosidase A (α-Gal A) activity. Classic Fabry disease affects around 1 in 40,000-60,000 males, leading to glycosphingolipid accumulation, primarily globotriaosylceramide (Gb3), causing childhood or adolescent symptoms like acroparesthesia, angiokeratoma, and hypohidrosis, progressing to renal insufficiency, cardiomyopathy, and cerebrovascular disease in adulthood. Later-onset phenotypes, with higher residual enzyme activity, present milder symptoms such as hypertrophic cardiomyopathy, renal failure, or cryptogenic stroke in adulthood. The cardiac phenotype manifests typically during the fifth to eighth decade with impairments like hypertrophic cardiomyopathy, mitral insufficiency, and arrhythmias. [1–5]

In Taiwan, our research team was the first to identify a remarkably high incidence of a later-onset cardiac type GLA mutation, IVS4 + 919G > A (c.640-801G>A), which affects approximately 1 in 1,600 males in our population [6]. Due to the high prevalence of IVS4 FD in Taiwan, which raised doubts about its pathogenicity, the Taiwan National Health Insurance Administration implemented a treatment guideline requiring IVS4 patients to undergo endomyocardial biopsies to confirm that their hypertrophic cardiomyopathy is indeed caused by FD before they can apply for enzyme replacement therapy (ERT) from the national health insurance. Therefore, our team started to conduct endomyocardial biopsies on these IVS4 patients. Until now, more than 100 patients with IVS4 mutation have received biopsies, and the results have shown typical histopathological changes, such as abundant diffuse cytoplasmic vacuolization (inclusion bodies) on hematoxylin and eosin (H&E) staining and lamellar myelin bodies in electron microscopy, in endomyocardiac biopsies of all ERT naïve IVS4 patients with left ventricular hypertrophy (LVH). This finding indicates that IVS4+919G>A mutation is indeed a real pathogenic mutation [7,8].

Tissue fibrosis is considered an irreversible event with limited therapeutic intervention available and is a negative prognostic factor for ERT [9]. It has demonstrated that initiating ERT for Fabry cardiomyopathy before the onset of myocardial fibrosis is crucial for achieving sustained improvement in myocardial morphology and function [10]. However, our current study has revealed that in IVS4 patients, even those without LVH, around 38.1% of males and 16.7% of females already had myocardial fibrosis in their hearts, as detected by late gadolinium enhancement heart MRI. In the same study, endomyocardial biopsies were performed on seventeen patients, and all of them had significant classical Fabry pathological changes (abundant inclusions and lamellar myelin bodies) in their endomyocardial biopsies [11].

Unexpectedly, we later discovered that three patients with an IVS4 mutation, who exhibited some symptoms/signs of Fabry cardiomyopathy but did not have LVH. Cardiac biopsies were performed, but no typical Fabry pathological changes, such as inclusion bodies by optical microscope or lamellar myelin bodies by electron microscope, were found. However, Gb3 immunostaining revealed substantial Gb3 accumulation in all of these biopsies, and even extralysosomal Gb3 could be found in one of their biopsies [12]. This finding raises an interesting issue to investigate whether this early-stage Gb3 accumulation without typical Fabry pathogenic findings (inclusion bodies) could initiate severe inflammatory changes or even irreversible damage to cellular or organ function in these Fabry disease patients.

In this study, we initially assessed the severity of inflammatory and oxidative stress in fibroblasts obtained from FD patients with the IVS4 mutation compared to non-FD controls. This evaluation involved examining levels of inducible nitric oxide synthase (iNOS), phospho-p44/42 mitogen-activated protein kinase (MAPK), and interleukin-18 (IL-18) through immunofluorescence staining to assess the degree of inflammation and oxidative stress and further compare the severity between these two groups. Additionally, we performed myocardial biopsies on crossbred G3Stg/GLAko symptomatic mice and wild-type C57BL/6 mice. We also included the myocardial biopsies of the three FD patients mentioned earlier, which exhibited substantial Gb3 accumulation through immunostaining but did not display typical FD pathological changes. These samples underwent similar experiments to gauge the extent of inflammation and oxidative stress between the groups. In addition to immunostaining, western blot analysis was employed for myocardial biopsies of G3Stg/GLAko Fabry symptomatic mice and wild-type mice. Lastly, we performed immunofluorescence staining of alpha-smooth muscle actin (α-SMA) in myocardial biopsies from both mice and patients to detect the presence of fibrosis. The aim of the present study is to determine whether irreversible damage, such as cardiac fibrosis, could exist before the occurrence of typical pathological changes associated with Fabry disease. This information is crucial in selecting an appropriate time for the early initiation of treatment.

## Methods

### Fibroblast culture

Wild-type skin fibroblasts (BCRC number: 08C0011) and skin fibroblasts obtained from IVS4 FD patients (collected from Taipei Veterans General Hospital, IRB number: VGHUST105-G7-6-1) were cultivated in Dulbecco’s modified Eagle’s medium (Thermo Fisher Scientific, Waltham, MA, USA) supplemented with 10% heat-inactivated fetal bovine serum, 1% sodium pyruvate, and 1% L-glutamine in a 100 mm dish. The cells were maintained at a temperature of 37 °C in an atmosphere containing 5% CO2, and the medium was refreshed every 72 hours. For the immunofluorescence assay, the cells were seeded in 6-well plates at a density of 2×10^5^ cells per well and incubated in Dulbecco’s modified Eagle’s medium containing 10% FBS for 24 hours, until they reached 80% confluency.

### G3Stg/GLAko mice preparation

In a previous study, it was observed that G3Stg/GLAko mice exhibited a higher level of Gb3 accumulation compared to GLAko mice. Furthermore, the symptoms displayed by G3Stg/GLAko mice more closely resembled those observed in humans with FD [13]. Therefore, we chose G3Stg/GLAko mice as our experimental model. Transgenic human Gb3 synthase (TgG3S) mice were prepared as previously described and maintained by breeding with wild-type C57BL/6 mice [14]. The G3Stg/GLAko mouse line was generated by crossbreeding male TgG3S mice with homozygous female GLAko mice [15]. We selected 37-week-old wild-type C57BL/6 and G3Stg/GLAko mice deliberately to mirror the age range of 30 to 40 years in humans, typically when initial cardiac symptoms appear in IVS4 FD patients. Additionally, hearts harvested from 37-week-old G3Stg/GLAko mice showed Gb3 accumulation via immunostaining but lacked typical FD-associated pathological changes (inclusion bodies), closely resembling those observed in our patient cohort. This observation underscores the suitability of this age group of mice for our study. All procedures followed the principles and guidelines outlined in the Guidelines on Laboratory Animal Use and Management established by the ROC Council of Agriculture. Our studies were approved by the Institutional Animal Care and Use Committee at Taipei Veterans General Hospital.

### Patient Biopsy Samples

Three male patients carrying an IVS4 mutation, displaying symptoms of Fabry cardiomyopathy but without LVH, had their previous cardiac biopsies retrieved for inflammatory and fibrosis immunostaining investigations. None of the patients had received ERT before the biopsy. Despite the absence of typical Fabry pathological features, Gb3 immunostaining revealed significant Gb3 accumulation in the biopsies [12]. The study protocol was approved by the institutional review boards of Taipei Veterans General Hospital (IRB number: VGHUST105-G7-6-1), and all participants provided written informed consent.

### Case 1

A 44-year-old male with an IVS4 FD mutation and a family history of Fabry cardiomyopathy presented with intermittent chest pain and tightness for over one year. While electrocardiography (ECG) showed complete right bundle branch block (RBBB), echocardiography revealed no evidence of left ventricular hypertrophy (left ventricular mass index (LVMI): 26.73 g/m^2.7; normal <51). Initial lab work indicated elevated serum lyso-Gb3 (2.78 nM; normal <2.0 nM) and reduced plasma α-galactosidase A activity (1.09 nmol/h/ml; normal: 7.9–16.9 nmol/h/ml). Cardiac MRI detected subendocardial late gadolinium enhancement in the inferior and anterolateral left ventricular segments. However, routine pathological examinations for Fabry disease on endomyocardial biopsy tissue showed no typical pathological changes of Fabry disease (inclusion bodies), but substantial Gb3 accumulation, including extralysosomal Gb3 accumulation, could be found by Gb3 immunostaining [12].

### Case 2

A 37-year-old male with IVS4 FD was diagnosed with a cryptogenic stroke. Neuroimaging revealed an acute pontine infarction and dissection with an intimal flap in the mid basilar artery. Additional findings included dolichoectasia of the vertebrobasilar vessels. While lab work showed decreased plasma α-galactosidase A activity (1.15 nmol/h/ml; normal: 7.9–16.9 nmol/h/ml) and elevated lyso-Gb3 (2.85 nM; normal <2.0 nM), the patient exhibited no significant signs of Fabry cardiomyopathy (ECG: normal, heart echo: LVMI: 46.85 g/m^2.7; normal <51). Endomyocardial biopsy was pursued for enzyme replacement therapy eligibility; however, routine pathological examination for Fabry disease showed no inclusion bodies, but substantial Gb3 accumulation could be found by Gb3 immunostaining [12].

### Case 3

A 41-year-old male with chest tightness and palpitations was diagnosed with IVS4 FD during family screening. Initial lab work revealed elevated lyso-Gb3 (3.75 nM; normal <2.0 nM) and reduced plasma enzyme activity (1.29 nmol/h/ml; normal: 7.9–16.9 nmol/h/ml). Cardiac evaluation, including ECG, MRI, and echocardiography, showed no evidence of Fabry cardiomyopathy (LVMI: 29.46 g/m^2.7; normal <51). Despite this, an endomyocardial biopsy was performed at the patient’s request. Routine pathological examinations for Fabry disease did not demonstrate typical pathological changes, but substantial Gb3 accumulation could be found by Gb3 immunostaining [12].

### Immunofluorescent staining and quantification of Gb3, IL-18, iNOS, phospho-p42/44 MAPK and α-SMA accumulations in fibroblast and myocardial biopsy

Multiple studies have implicated inflammatory markers, including IL-18 and p42/44 MAPK, in the development of Fabry cardiomyopathy [16–19]. Additionally, increased levels of iNOS have been found to correlate with cardiac dysfunction in FD and are even associated with cardiomyocyte death [20]. Furthermore, α-SMA has been identified as definitive markers of cardiac fibrosis [21,22]. In this study, we aimed to investigate whether cellular stress, inflammation, and irreversible damage could already exist in the early stages of Gb3 accumulation in Fabry disease by examining these biomarkers.

For our experimental analysis, we utilized fibroblasts obtained from individuals diagnosed with FD carrying the IVS4 mutation, as well as non-FD controls. Additionally, myocardial biopsies from G3Stg/GLAko mice, wild-type mice, and FD patients with the IVS4 mutation were included. Following the removal of the growth medium, the cells were washed with phosphate-buffered saline (PBS) to eliminate any residual culture media. Subsequently, the cells were fixed using Fixation/Permeabilization solution (BD) for 20 minutes at room temperature. To block non-specific binding sites, the cells were incubated with 5% Fetal Bovine Serum (FBS) for 1 hour at room temperature.

Primary antibodies listed in **Table 1** were applied to the cells overnight at 4°C. After incubation, the cells were washed with 1X Perm/wash solution (BD) to remove any unbound primary antibodies. The cells were then incubated with the corresponding fluorescently labeled secondary antibodies listed in **Table 1** for 2 hours at room temperature. Nuclear staining was performed using 4,6-diamidino-2-phenylindole (DAPI), while lysosomal staining in the Gb3 group utilized lysosomal-associated membrane protein 1 (LAMP-1) as the lysosomal marker. Excess liquid was removed, and the stained cells were mounted on glass slides using Fluorescence-Mounting solution (Dako Omnis, Santa Clara, CA, USA).

**Table 1.** Primary and secondary antibodies of immunofluorescent staining.

| Antibodies | Host | Dilution | Structures<br>Identified by<br>Antibodies | Source |
| --- | --- | --- | --- | --- |
| 18985-1-AP<br>( polyclonal antibody) | Rabbit | 1:200 | iNOS | Proteintech<br>(Rosemont, IL, USA) |
| 10745-1-AP<br>(polyclonal antibody) | Rabbit | 1:200 | Nf-kB | Proteintech<br>(Rosemont, IL, USA) |
| 10663-1-AP<br>(polyclonal antibody) | Rabbit | 1:200 | IL-18 | Proteintech<br>(Rosemont, IL, USA) |
| C6198<br>(monoclonal antibody) | Rabbit | 1:400 | α-SMA | Sigma<br>( St. Louis , MO, USA) |
| A2506<br>(Monoclonal antibody) | Mouse | 1:200 | Gb3 | Tokyo Chemical Industry<br>(Tokyo, Japan) |
| #4370T<br>( polyclonal antibody) | Rabbit | 1:2000 | p-p42/44 | Cell Signaling Technology<br>(Danvers, MA, USA) |
| Dylight488 Goat<br>GTX213111-04 | Mouse | 1:1000 | Secondary<br>antibody | Genetex<br>(Irvine , CA, USA) |
| Dylight594 Goat<br>GTX213110-05 | Rabbit | 1:1000 | Secondary<br>antibody | Genetex<br>(Irvine , CA, USA) |

Finally, the mounted sections were examined under a fluorescence microscope (Nikon ECLIPSE Ni-E, Tokyo, Japan) equipped with a DP71 digital camera (Olympus) at a magnification of x200 and x600. The acquired images were processed and assembled using Image J software (National Institutes of Health, Bethesda, MD, USA) to facilitate analysis and comparison.

To quantify the levels of Gb3, IL-18, and iNOS in IVS4 fibroblasts and cardiomyocytes from patients, in G3Stg/GLAko mice, as well as in normal control samples, we utilized image analysis techniques. Specifically, we utilized ImageJ software (National Institutes of Health, Bethesda, MD, USA) to calculate the corrected total cell fluorescence (CTCF) of each section. This approach allowed us to determine the fluorescence intensity of the target protein and normalize it to the number of nuclei present in the sample. For each individual sample, a minimum of two sets of image fields containing approximately 100 cells were acquired, averaged, and employed for the computation of CTCF.

### Western blot and quantification of iNOS, IL-18 and phospho-p42/44 MAPK in mouse myocardial biopsy

Myocardial samples from G3Stg/GLAko mice and wild-type mice were prepared in SDS buffer (125mM Tris-HCl pH 6.8, 5% glycerol, 4% SDS, 0.005% bromophenol blue, 10% β-mercaptoethanol) and separated by 12% SDS-PAGE. Proteins were transferred to polyvinylidene fluoride (PVDF) membranes (Immobilon-P, Merck Millipore), blocked with 5% non-fat dry milk, and incubated with primary antibodies for 1h at room temperature. Membranes were then incubated for 45min at room temperature with horseradish peroxidase-conjugated secondary antibodies (anti-rabbit IgG). Protein bands were visualized using enhanced chemiluminescence (PerkinElmer) and imaged with an ImageQuant LAS 4000 (GE Healthcare). Band densities were quantified by Multi Gauge software (Fujifilm) for densitometric analyses. The primary and secondary antibodies utilized for Western blotting are listed in **Table 2**.

**Table 2.** Primary and secondary antibodies of Western blot.

| Antibodies | Host | Dilution | Structures<br>Identified by<br>Antibodies | Source |
| --- | --- | --- | --- | --- |
| 18985-1-AP<br>(polyclonal antibody) | Rabbit | 1:1000 | iNOS | Proteintech<br>(Rosemont, IL, USA) |
| 10663-1-AP<br>(polyclonal antibody) | Rabbit | 1:3000 | IL-18 | Proteintech<br>(Rosemont, IL, USA) |
| #4370T<br>( polyclonal antibody) | Rabbit | 1:2000 | p-p42/44 | Cell Signaling Technology<br>(Danvers, MA, USA) |
| #4695T<br>( polyclonal antibody) | Rabbit | 1:2000 | p42/44 | Cell Signaling Technology<br>(Danvers, MA, USA) |
| GTX110564 | Rabbit | 1:6000 | β-actin | Genetex<br>(Irvine , CA, USA) |
| GTX100118 | Rabbit | 1:6000 | GAPDH | Genetex<br>(Irvine , CA, USA) |

The relative abundance of target proteins from Western blotting was determined by densitometric analysis using ImageJ software (NIH). Digitized images of chemiluminescent Western blot band signals were converted to 8-bit grayscale and inverted. Using the manual band selection tool, rectangle areas of identical sizes were drawn around each protein band to measure density in pixel intensity units. Background subtraction was applied by selecting a region without bands near each lane. The resulting band densities were normalized to that of the housekeeping protein GAPDH and β-actin in the same lane to account for any variations in total protein loading across lanes. Normalized band densities were then expressed as fold changes relative to the control sample band density set at 1.

All statistical analysis of immunofluorescent staining and Western blot were performed using MedCalc version 22.006 (MedCalc Software Ltd., Ostend, Belgium). The Mann-Whitney U test and Kruskal-Wallis test were employed to evaluate the differences between groups due to the small sample size. Statistical significance was determined with a *p*-value threshold of 0.05, indicating the presence of a significant difference between the compared groups.

## Results

### Early Gb3 Accumulation Correlates with Elevated Inflammatory Biomarkers in Fabry *Disease Fibroblasts*

To further investigate the presence of inflammation, we conducted a comparative analysis of inflammatory biomarkers in normal human and Fabry disease-derived fibroblasts. Initially, we confirmed the accumulation of globotriaosylceramide (Gb3) in Fabry disease-derived fibroblasts before quantifying the expression levels of biomarkers. Our analysis revealed a significant increase in Gb3 levels in the IVS4 Fabry disease (FD) fibroblasts compared to the control group (p < 0.001) (Figure 1a, b). Notably, similar elevations were observed for inducible nitric oxide synthase (iNOS) (p < 0.001) (Figure 1c, d), interleukin-18 (IL-18) (p < 0.001) (Figure 1e, f), nuclear factor kappa B (NF-κB)(p < 0.001) (Figure 1g, h), phosphorylated p42/44 mitogen-activated protein kinase (phospho-p42/44 MAPK) (p = 0.0015) (Figure 1i, j), and alpha-smooth muscle actin (α-SMA) (p = 0.0057) (Figure 1k, l). These findings demonstrate a clear correlation between IVS4 FD and markedly elevated levels of Gb3 accumulation, concurrent with increased activation of the IL-18, iNOS, phospho-p42/44 MAPK, and α-SMA signaling pathways. Given that Gb3 accumulation in these Fabry disease-derived fibroblasts is detectable only through Gb3 immunostaining, and classical Fabry pathological changes are not identifiable through routine pathological examination, we posit that the Gb3 accumulation in these fibroblasts represents a very early stage of the disease process. Consequently, these data lend further support to our hypothesis that early Gb3 accumulation initiates an inflammatory response.

**Figure 1.**
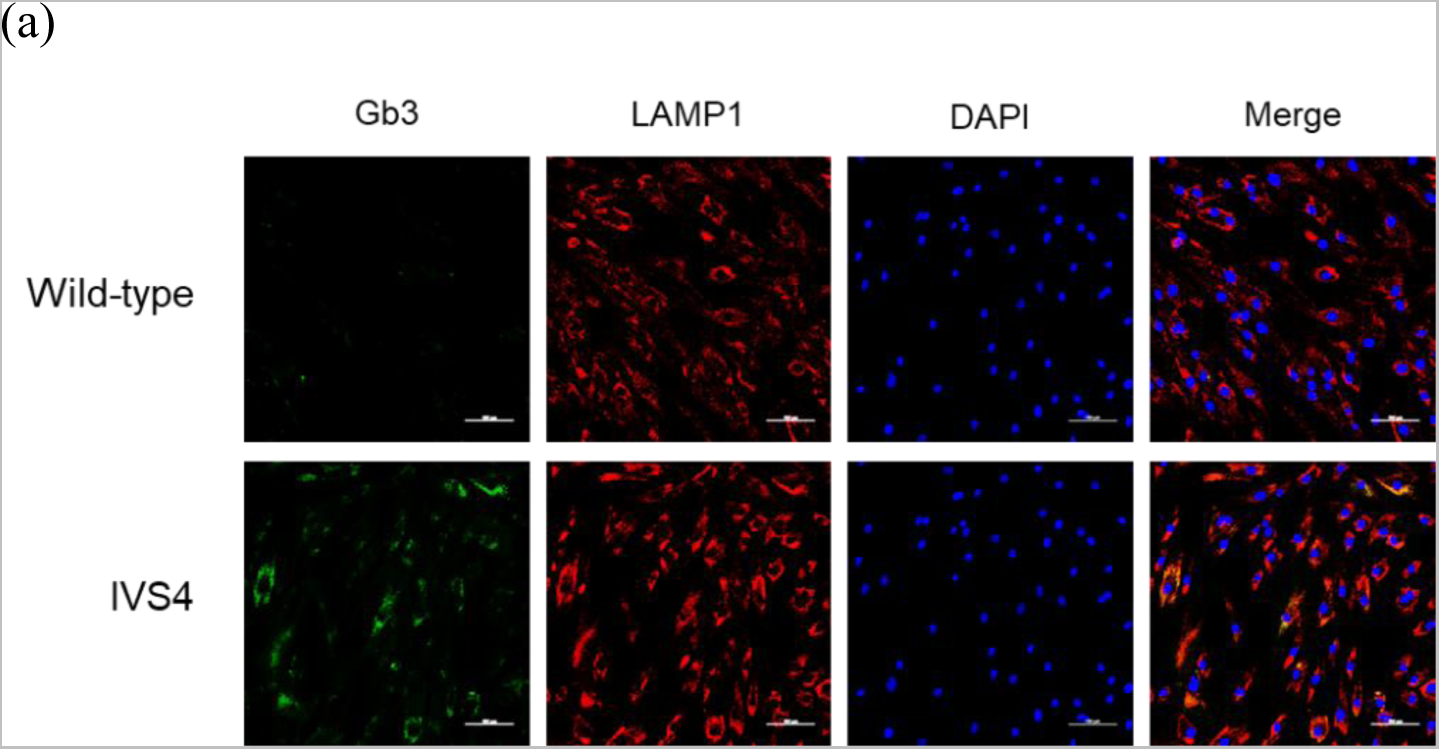
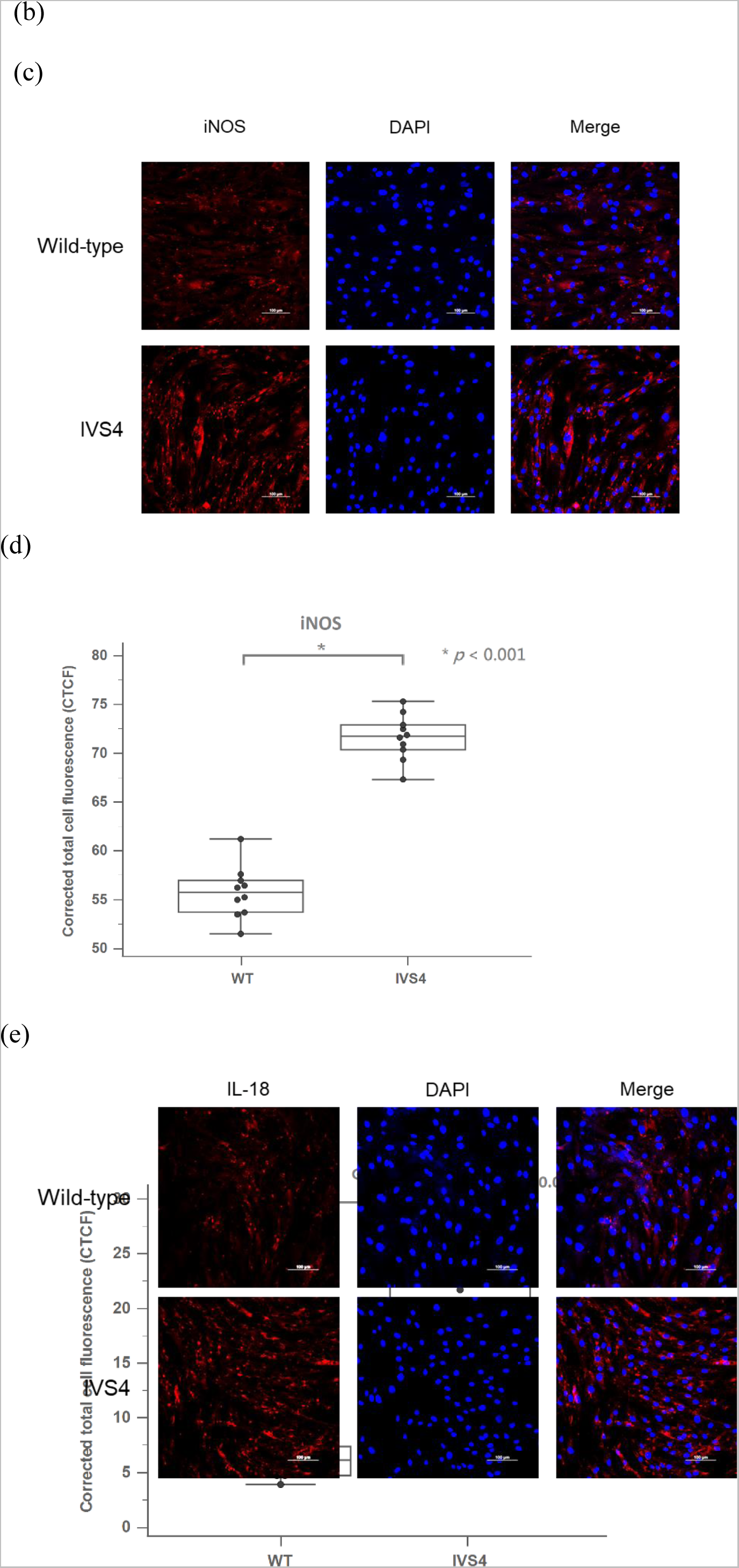
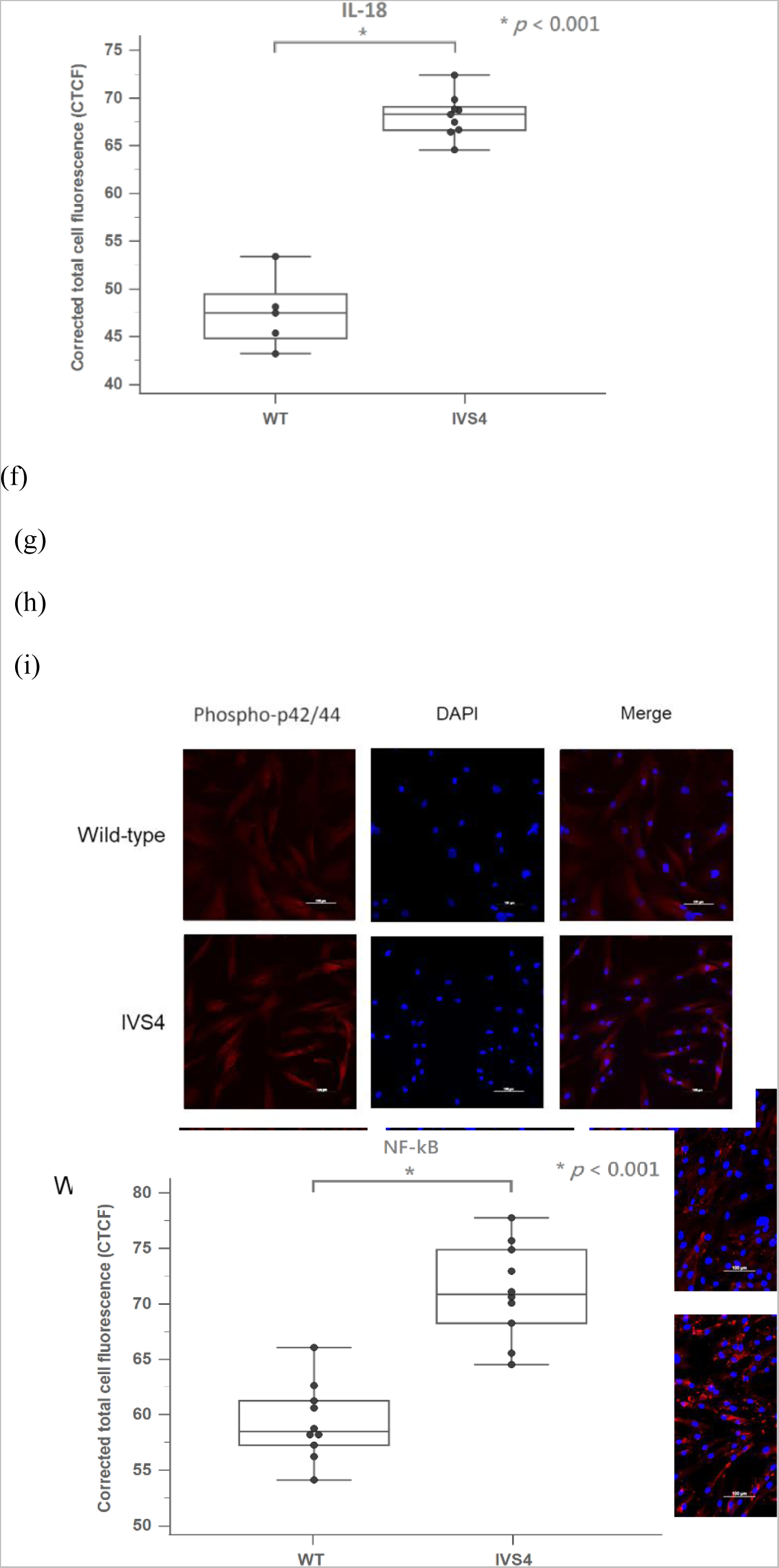
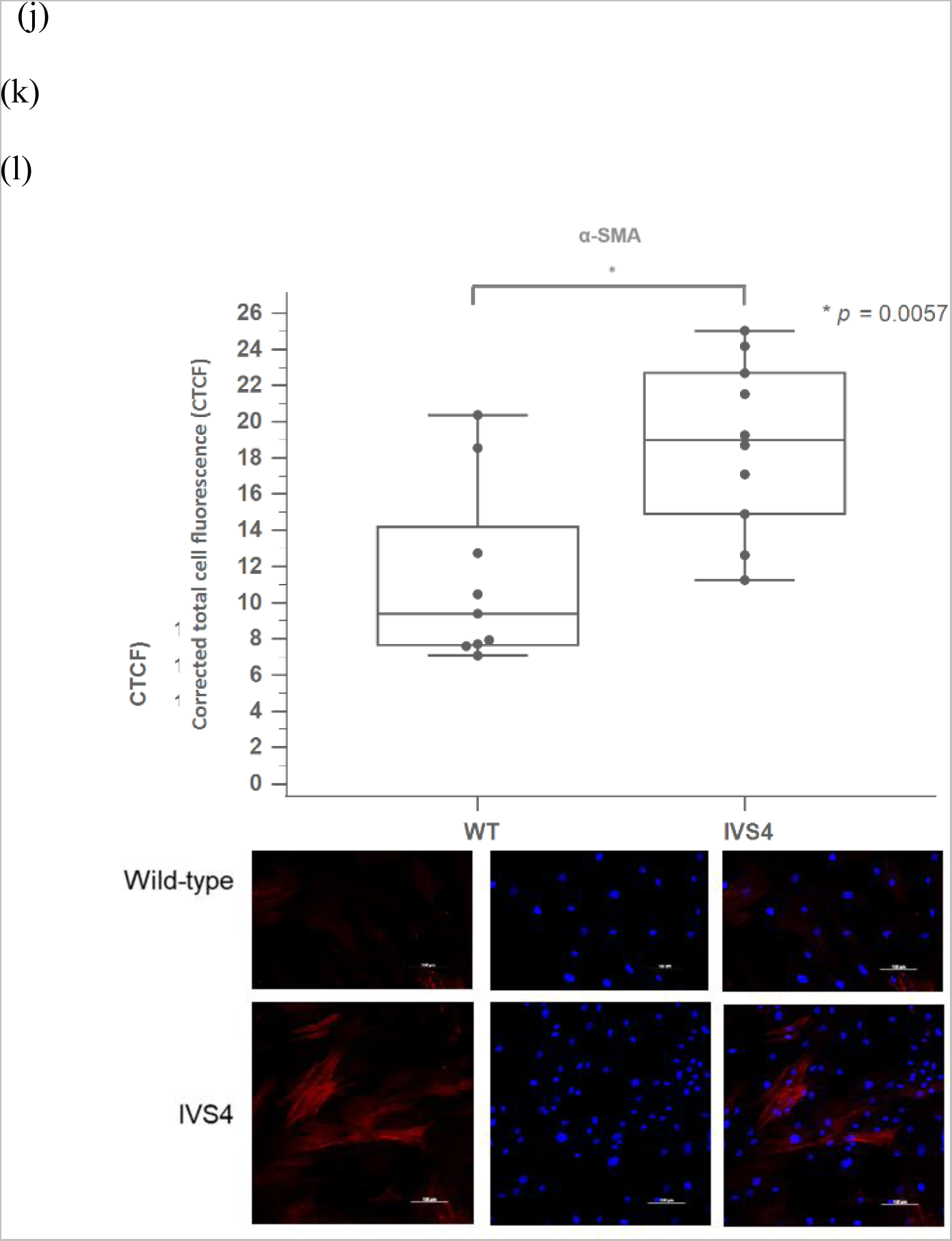
Immunofluorescence staining and quantification of Gb3 and inflammatory biomarkers in fibroblasts. (a) Immunofluorescence staining of Gb3 (green) and LAMP-1 (red), a lysosomal marker, in control and IVS4 Fabry disease (FD) fibroblasts. (b) Quantification of corrected total cell fluorescence (CTCF) for Gb3. (c, d) Immunofluorescence staining and CTCF quantification for iNOS, an oxidative stress marker. (e, f, g, h) Immunofluorescence staining and CTCF quantification for IL-18 and NF-κB, both are inflammatory markers. (i, j) Immunofluorescence staining and CTCF quantification for phospho-p42/44 MAPK, a signaling molecule involved in inflammation. (k, l) Immunofluorescence staining and CTCF quantification for α-SMA, a marker of myofibroblasts and fibrosis. DAPI (blue) was used for nuclear counterstaining. Data are presented as mean ± SEM. Scale bar = 100 μm.

### Confirmation of Inflammatory and Fibrotic Markers in a Fabry Disease Mouse Model and Human Cardiac Biopsies

Due to the challenges in obtaining cardiac tissue samples from human normal controls and the limitations of fibroblast models in fully representing the disease phenotype, we utilized a previously published symptomatic Fabry disease mouse model (G3Stg/GLAko) and 3 IVS4 patients, and compared them to a wild-type mouse model to confirm the elevation and presence of inflammation and fibrosis in myocardial biopsies of the symptomatic mouse model and IVS4 patients. The results of globotriaosylceramide (Gb3) immunofluorescence analysis for myocardial biopsies from three IVS4 patients have been previously reported [12]. These images clearly demonstrate that Gb3 accumulation in myocardial biopsies of G3Stg/GLAko mice and IVS4 patients is significantly higher compared to that in wild-type mice (Figure 2a).

**Figure 2.**
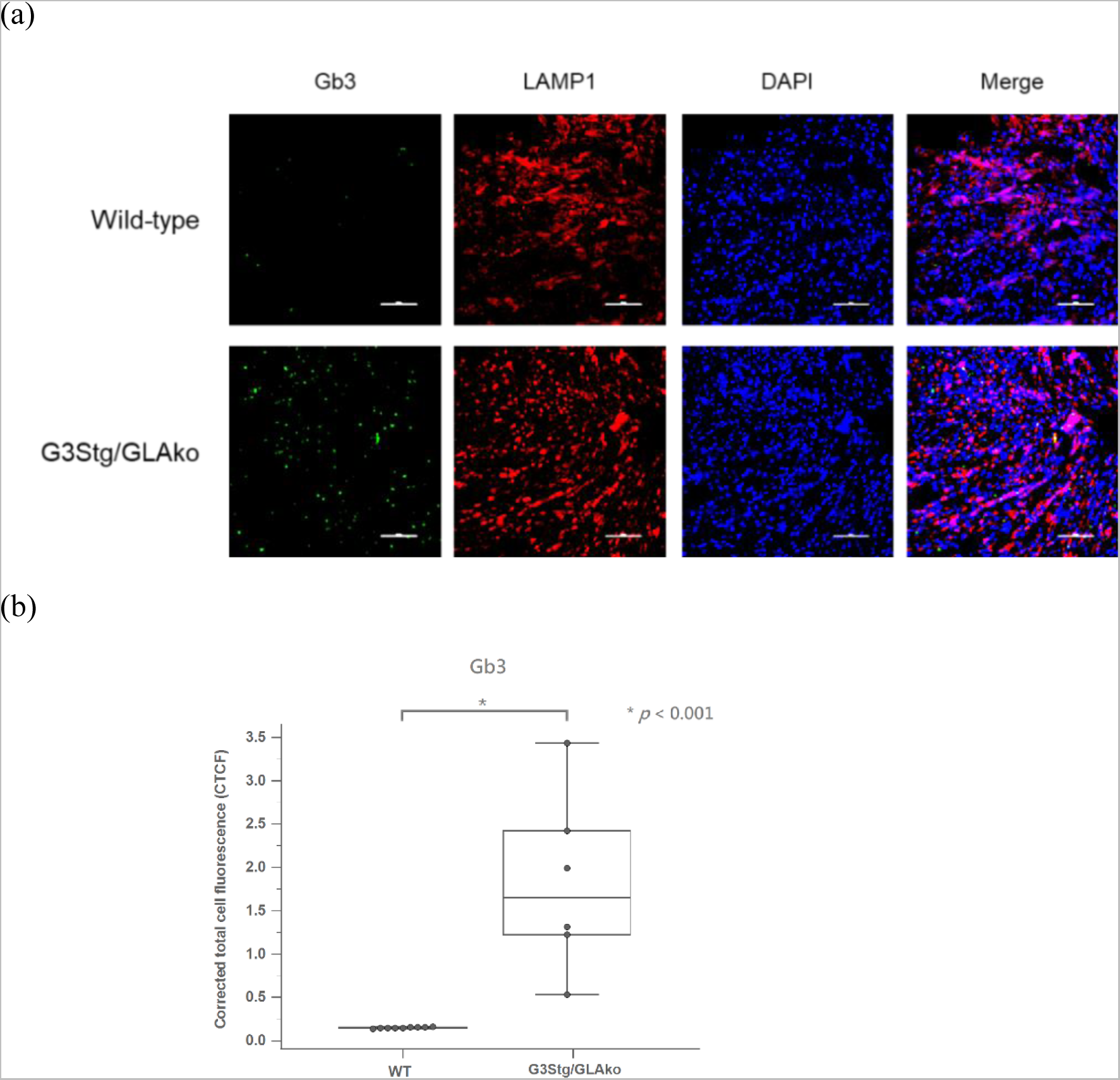
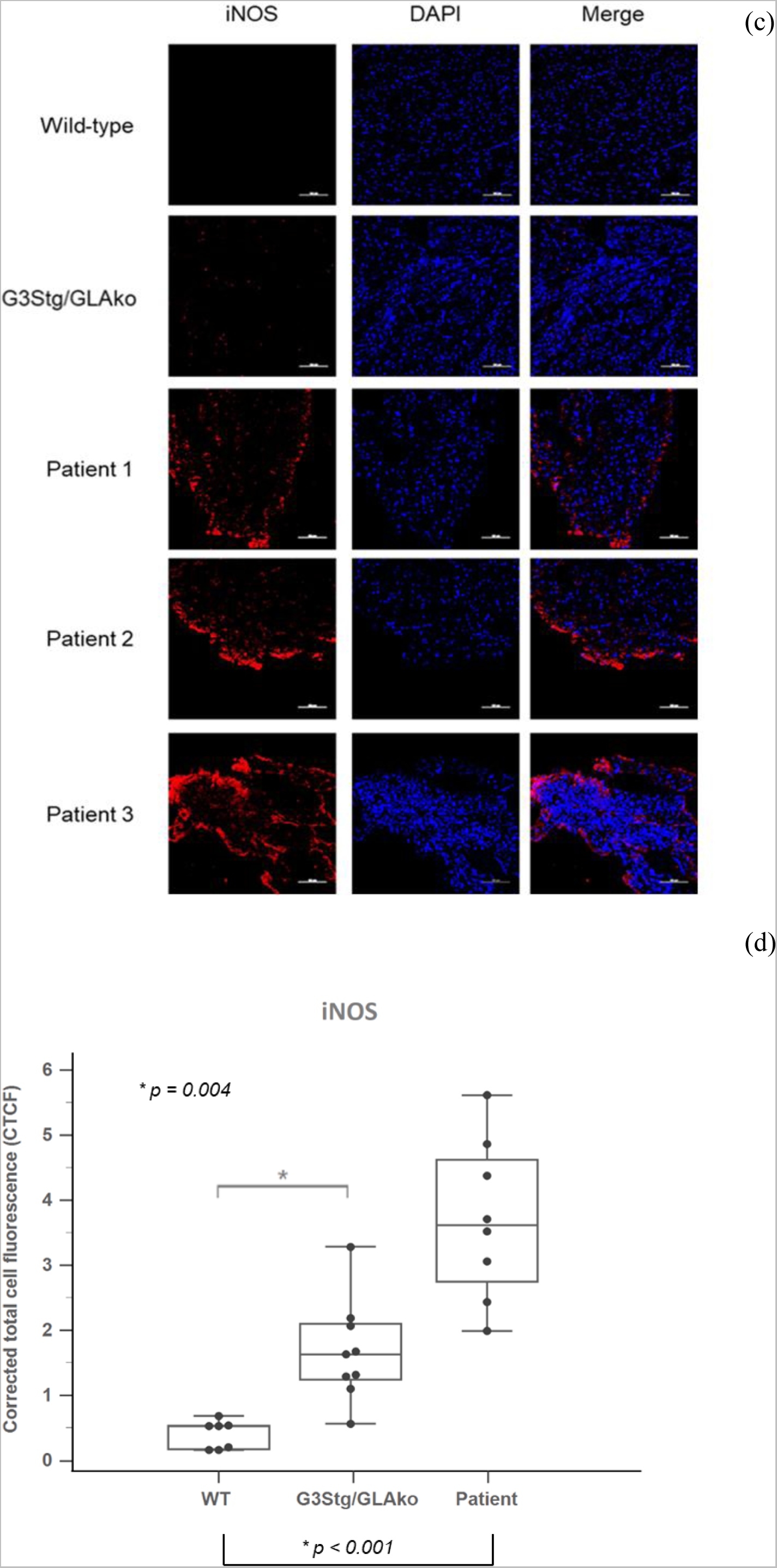
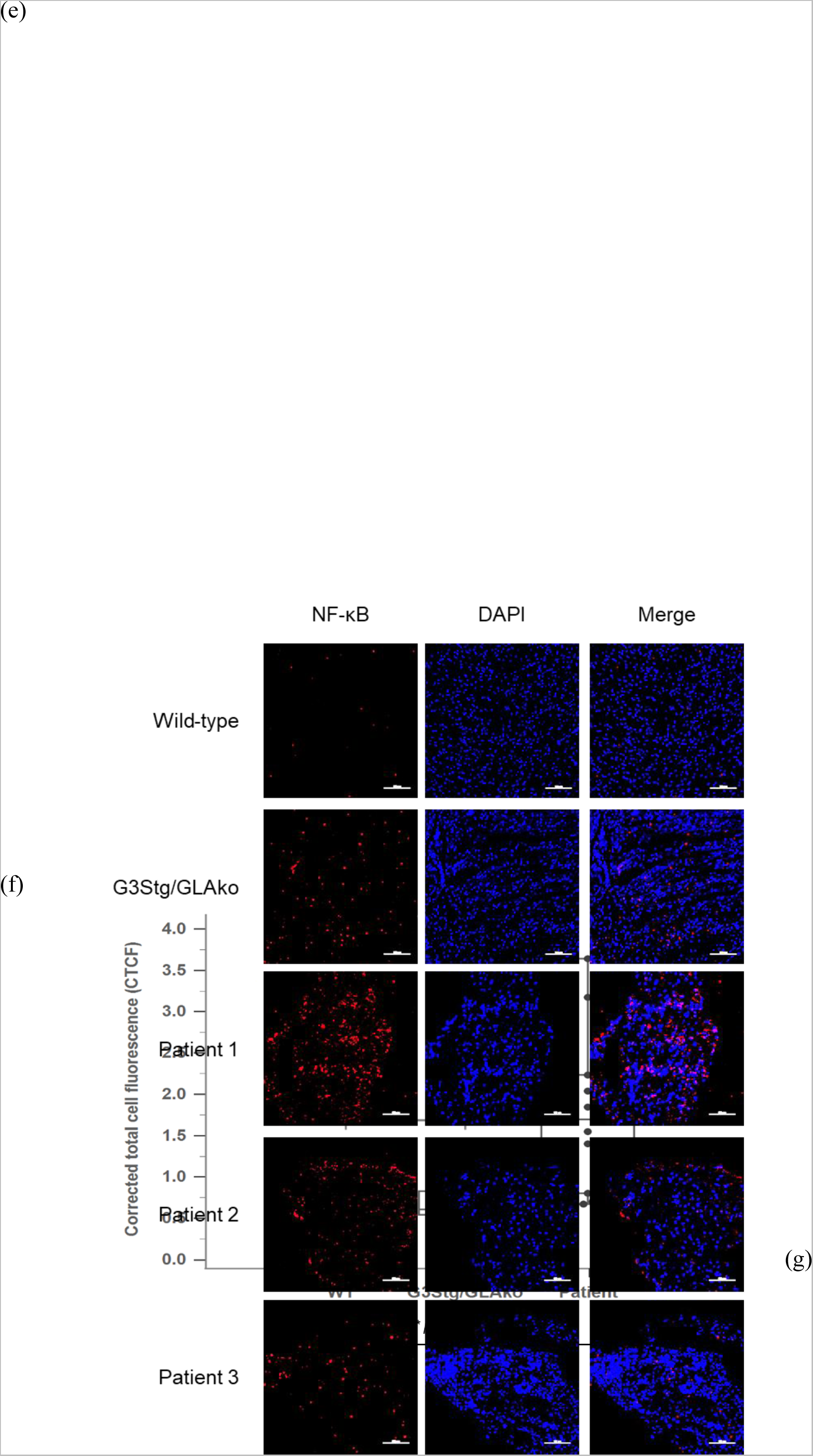
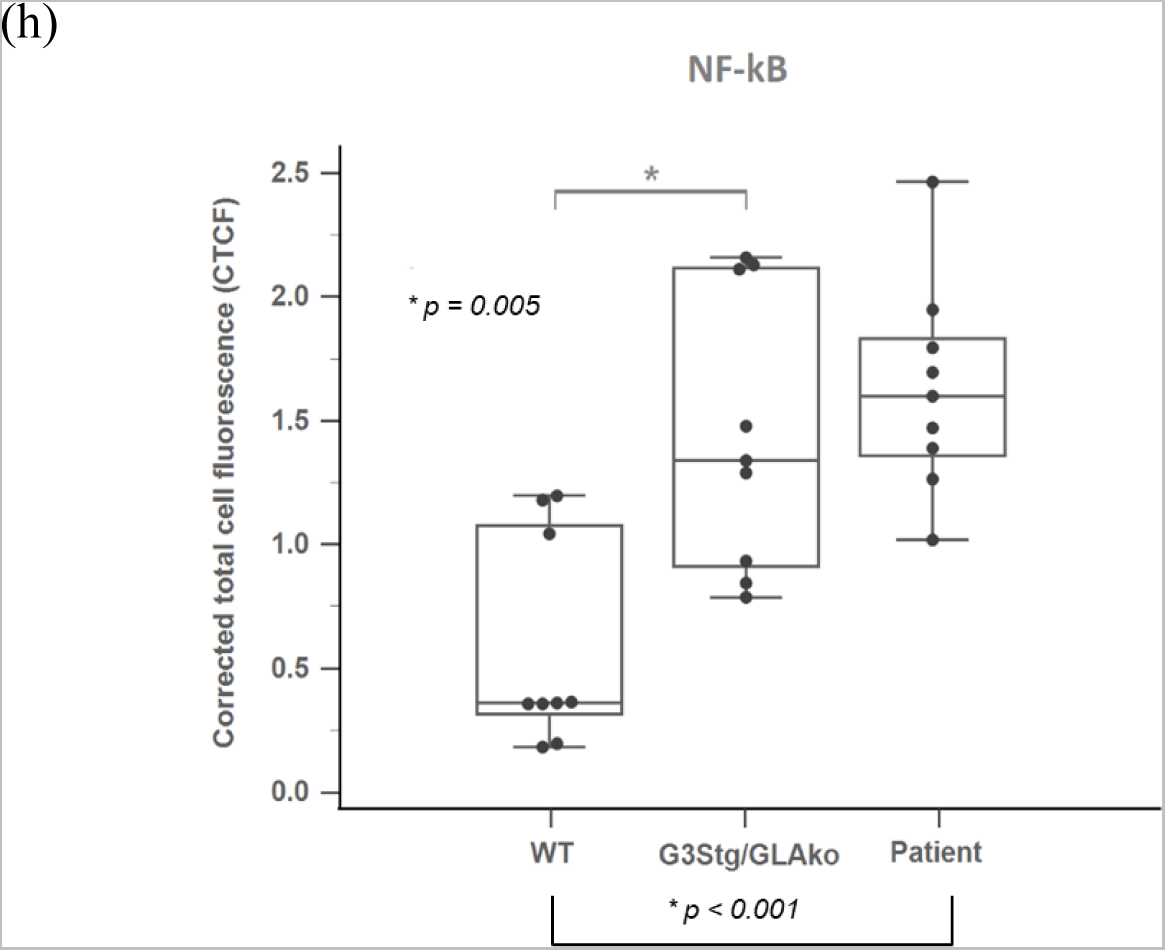
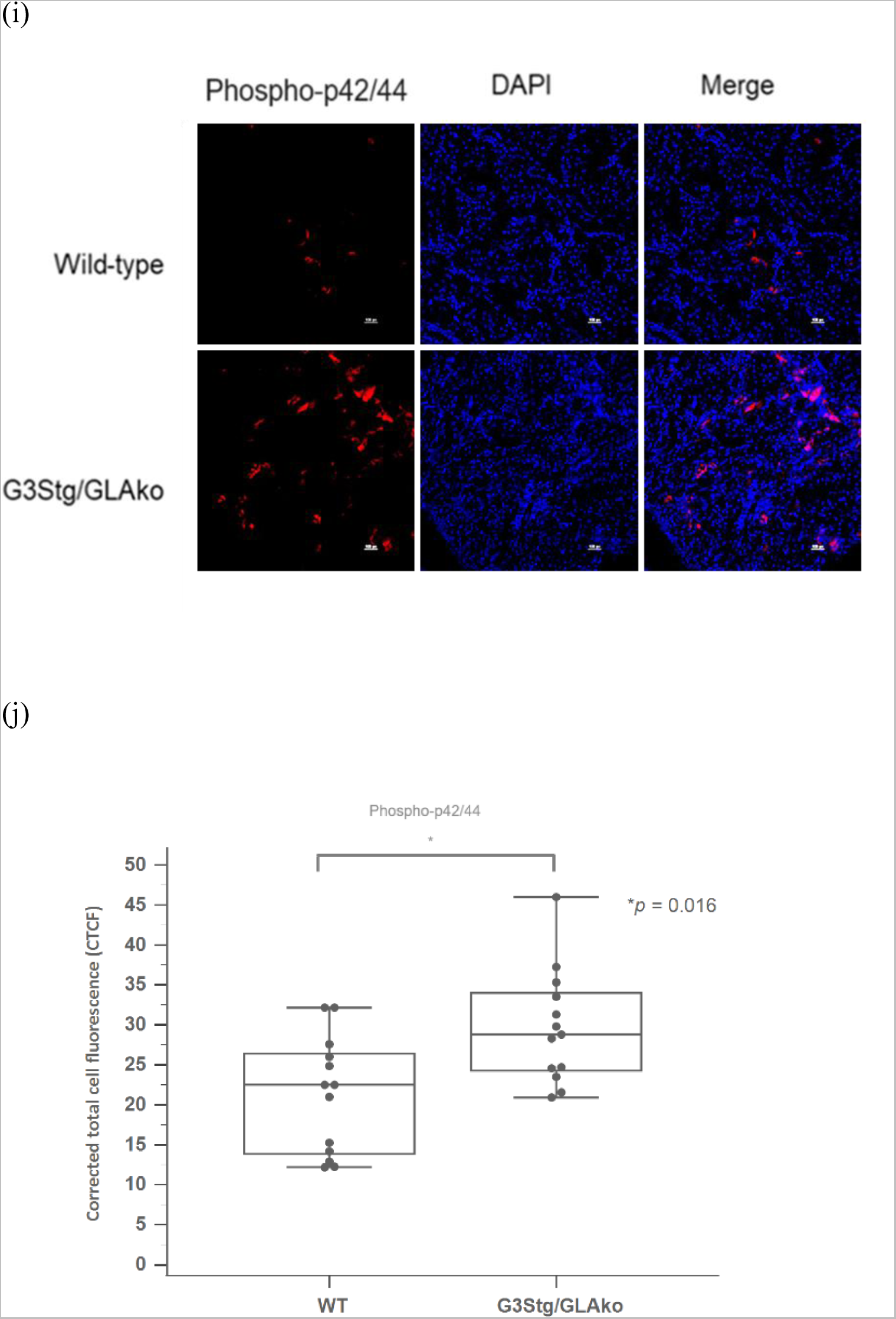
Analysis of Gb3 accumulation and inflammatory biomarkers in G3Stg/GLAko mice. (a) Immunofluorescence staining of Gb3 (green) and LAMP-1 (red) in myocardial sections from wild-type and G3Stg/GLAko mice, a Fabry disease mouse model. (b) Quantification of CTCF for Gb3. (c, d) Immunofluorescence staining and CTCF quantification for iNOS in myocardial sections from IVS4 FD patients, wild-type, and G3Stg/GLAko mice. (e, f, g, h) Immunofluorescence staining and CTCF quantification for IL-18 and NF-κB in myocardial sections from IVS4 FD patients, wild-type, and G3Stg/GLAko mice. (i, j) Immunofluorescence staining and CTCF quantification for phospho-p42/44 MAPK from wild-type and G3Stg/GLAko mice. DAPI (blue) was used for nuclear counterstaining. Data are presented as mean ± SEM. Scale bar = 100 μm.

We quantified the fluorescence intensity of cardiac biopsies from G3Stg/GLAko and wild-type mice. The fluorescence intensity of Gb3 in myocardial biopsies obtained from G3Stg/GLAko mice and IVS4 Fabry disease (FD) patients was significantly higher compared to that in wild-type mice (Figure 2b) (p < 0.001). Furthermore, we analyzed the expression of inflammatory biomarkers in myocardial biopsies of the mouse models and IVS4 patients using immunofluorescence staining. As shown in Figure 2, immunofluorescence staining for iNOS (Figure 2c), IL-18 (Figure 2e), NF-κB(Figure 2g), and phospho-p42/44 MAPK(Figure 2i) revealed elevated expression levels in the diseased mice and IVS4 FD patients, with statistically significant differences across all measured parameters (highest p-value = 0.009, other p-values were at 0.004 and 0.005 or even below 0.001)(Figure 2d, f, h, j).

### Western Blot Confirmation of Early Inflammatory Response and Oxidative Stress in Fabry Disease Mouse Model

Western blot analysis was conducted to further validate that early globotriaosylceramide (Gb3) accumulation could elicit a significant inflammatory response and oxidative stress in cardiac tissue. Due to the limited quantity of myocardial biopsy samples from IVS4 Fabry disease (FD) patients, Western blot analysis was performed exclusively on mouse biopsies. Immunoblotting for iNOS (Figure 3a) and IL-18 (Figure 3b) demonstrated consistent findings with increased expression in the disease model compared to controls. Additionally, Western blot analysis revealed elevated levels of phospho-p42/44 MAPK relative to wild-type mice (Figure 3b). Statistical analysis of the Western blot results, presented in Figure 3c, showed significant differences between the disease model and controls for all measured proteins, with p-values < 0.05. These results provide compelling evidence that prior to observable Gb3 inclusion body formation, G3Stg/GLAko mice already exhibit significantly heightened myocardial expression of pro-inflammatory mediators iNOS and IL-18, as well as enhanced MAPK signaling, further supporting the early onset of inflammatory processes in Fabry disease.

**Figure 3.**
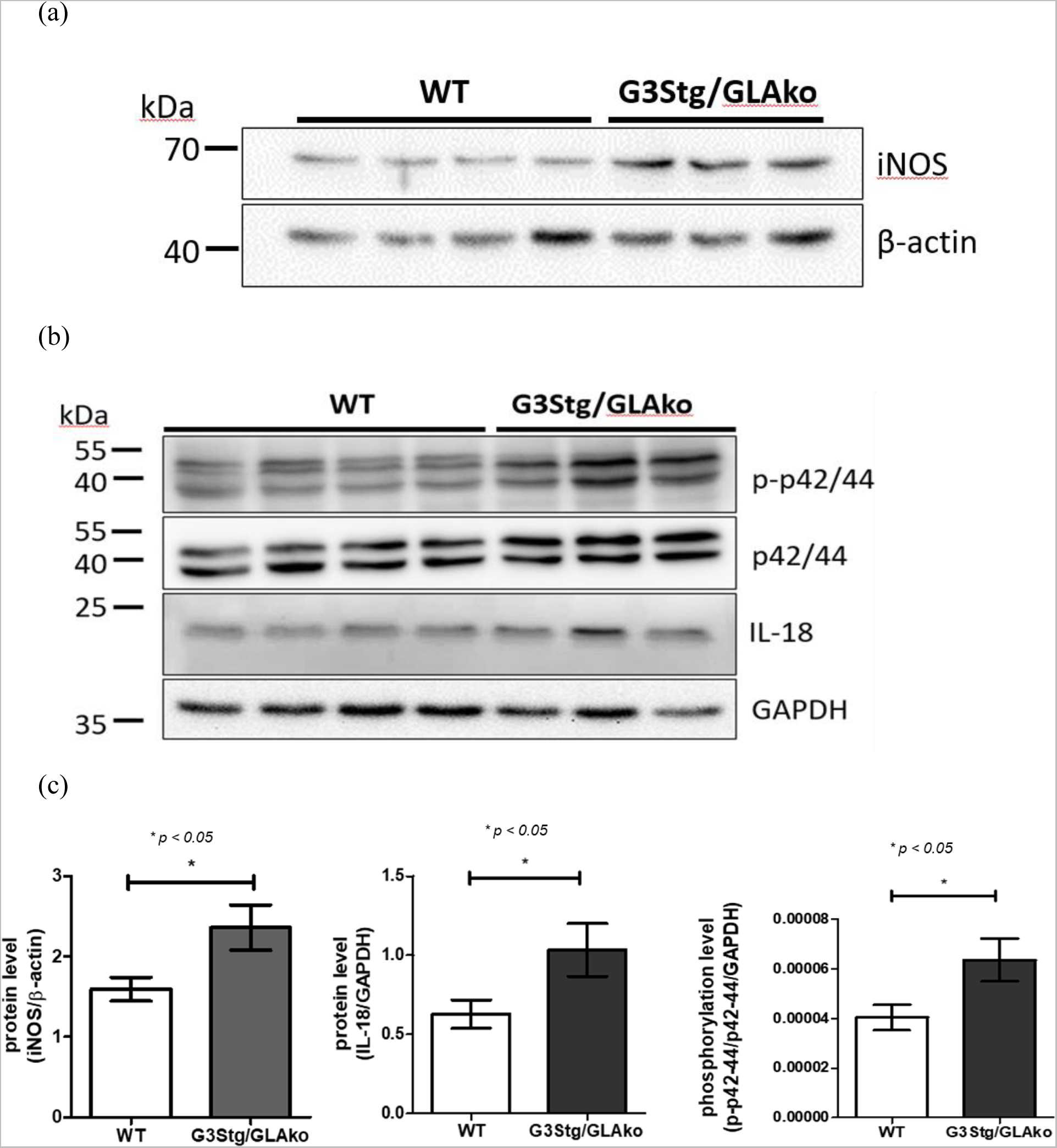
Western blot analysis of inflammatory markers and MAPK signaling in mouse myocardial biopsies. (a, b) Representative Western blot images for iNOS, IL-18, phospho-p42/44 MAPK, and loading controls (β-actin and GAPDH) in myocardial samples from wild-type and G3Stg/GLAko mice. (c) Quantification of relative protein levels for iNOS, IL-18, and phospho-p42/44 MAPK, respectively, normalized to loading controls. Data are presented as mean ± SEM. * *p* <0.05 compared to wild-type.

### Detection of Early-Stage Fibrosis in IVS4 Fabry Disease Patients and a Symptomatic Mouse Model

Based on the aforementioned data, we conducted further investigations to determine whether irreversible damage, such as inflammation-induced fibrosis, had already occurred in patients and the Fabry disease symptomatic mouse model. Alpha-smooth muscle actin (α-SMA) is a widely used immunohistochemical marker for detecting activated myofibroblasts and fibrosis [23]. When assessing fibrosis through α-SMA immunofluorescent staining on myocardial biopsies, it is crucial to consider the presence of blood vessels, such as capillaries, within the tissue sections, as they could potentially influence result interpretation. These vascular structures can be identified during microscopic examination, allowing the exclusion of their immunofluorescent staining signal from the analysis. Conversely, Western blot analysis for α-SMA typically requires a larger tissue sample, making it challenging to isolate pure populations of cardiomyocytes and endothelial cells from the biopsies. Therefore, Western blot analysis for α-SMA was not employed in this study.

Figure 4a illustrates the results of α-SMA immunofluorescence in myocardial biopsies from IVS4 Fabry disease (FD) patients. α-SMA expression was positive in all IVS4 FD patients, with notably higher expression observed in patient 1, who exhibited a more severe phenotype as previously described. These results suggest the existence of early-stage fibrosis in cardiomyocytes of IVS4 FD patients with globotriaosylceramide (Gb3) accumulation. We further evaluated the expression level of α-SMA in a Fabry disease symptomatic mouse model. The results of α-SMA immunofluorescence comparing myocardial biopsies in G3Stg/GLAko mice and wild-type mice are presented in Figure 4a. α-SMA expression was significantly higher in G3Stg/GLAko mice (p = 0.032) (Figure 4b). By integrating data from cardiac tissue samples of IVS4 FD patients and the mouse model, our findings indicate the presence of fibrosis in the early stages of Gb3 accumulation in cardiomyocytes in both IVS4 FD patients and G3Stg/GLAko mice.

**Figure 4.**
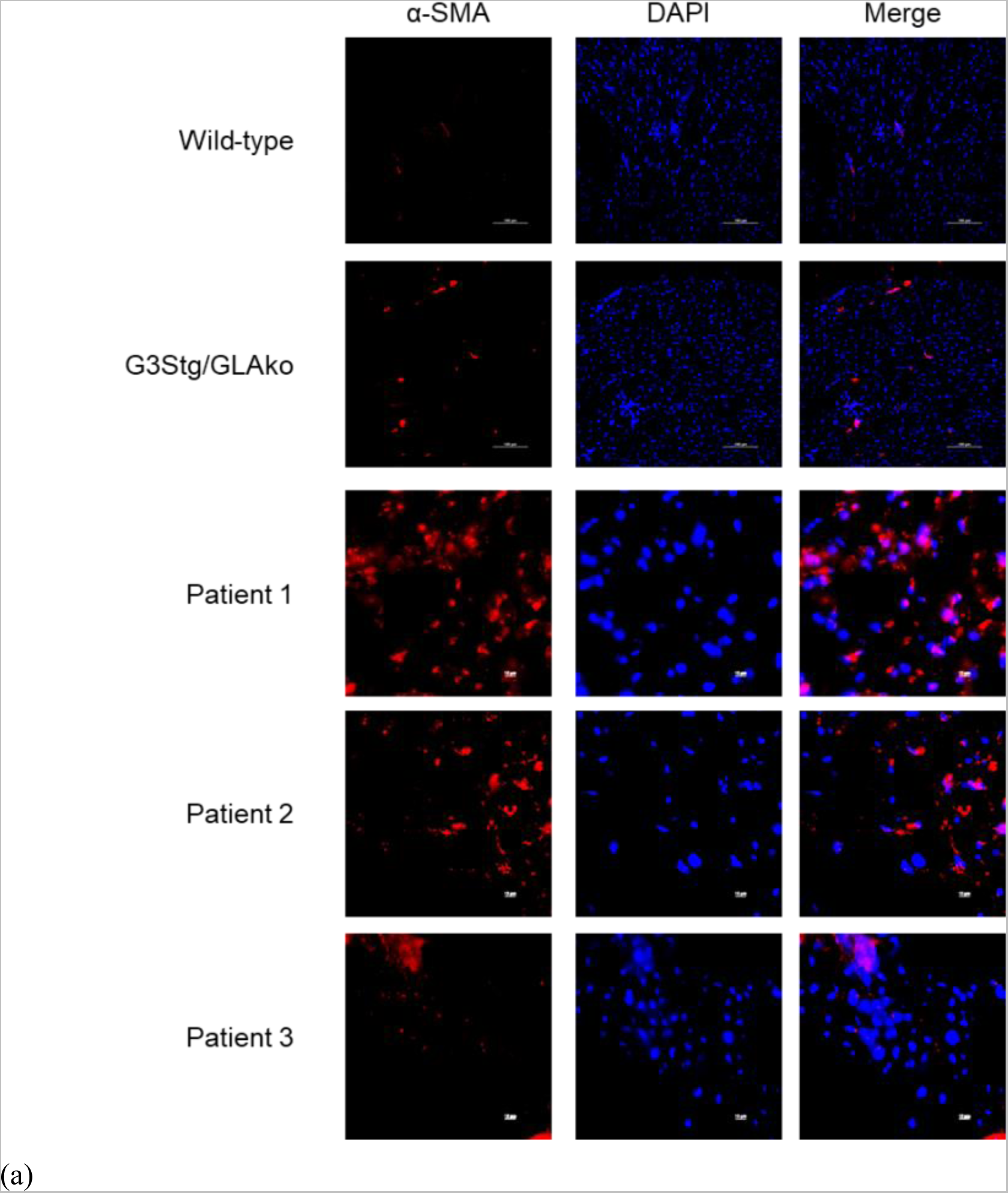
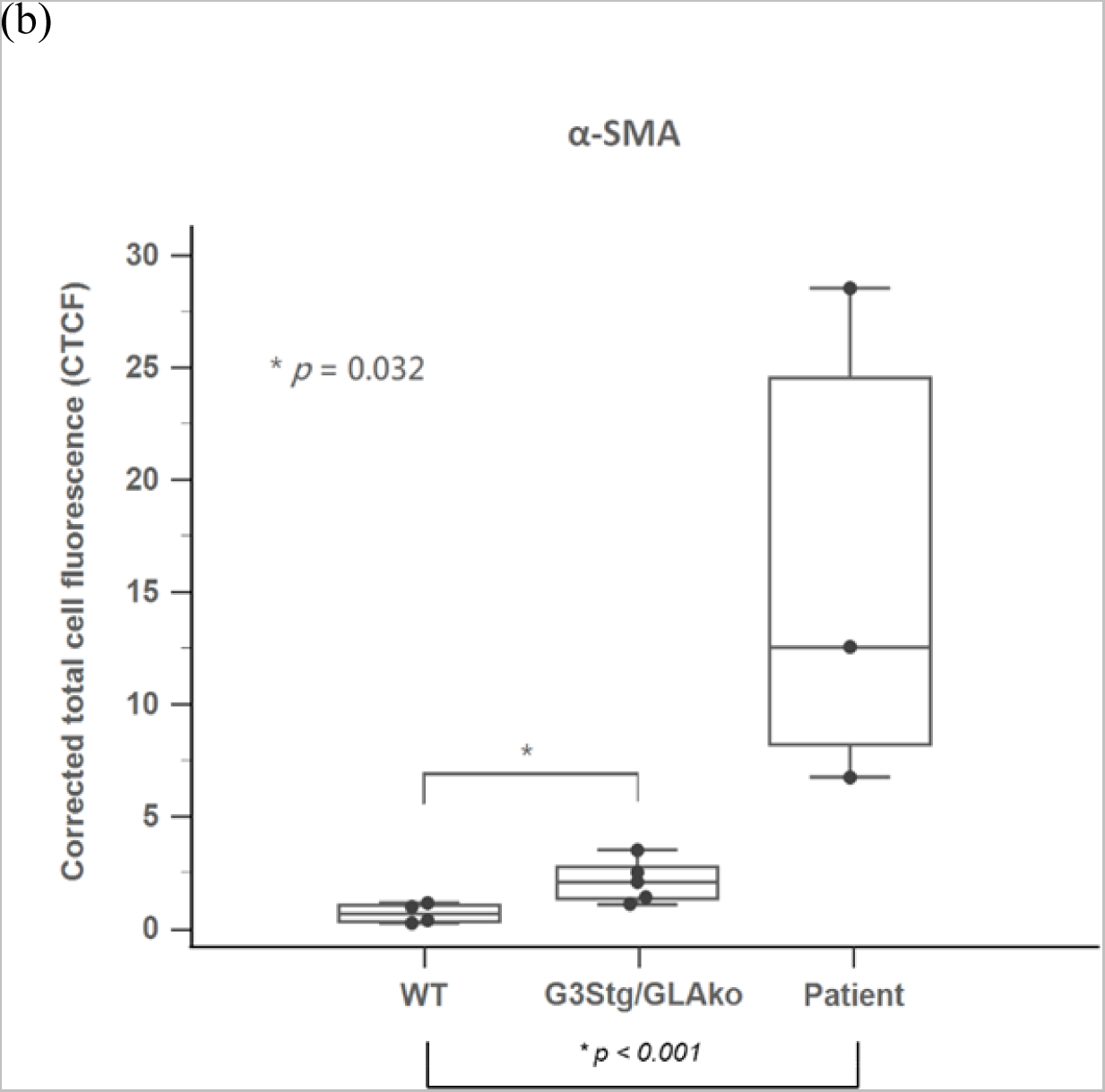
Immunofluorescence staining of α-SMA in cardiac biopsies. (a) Representative images of α-SMA staining (red) in myocardial sections from IVS4 FD patients, wild-type and G3Stg/GLAko mice. (b) Quantification of CTCF for α-SMA in myocardial sections from IVS4 FD patients, wild-type, and G3Stg/GLAko mice. DAPI (blue) was used for nuclear counterstaining. Data are presented as mean ± SEM. Scale bar = 100 μm.

## Discussion

The findings of this study provide important insights into the early-stage accumulation of Gb3 in FD patients and its impact on cardiomyocytes. Traditionally, the presence of Gb3 inclusion bodies has been considered the hallmark pathological change in FD cardiomyopathy [5]. However, this study challenges this notion by demonstrating that significant cellular stress and irreversible damage may already exist before the typical appearance of inclusion bodies.

The study showed that cardiomyocytes without inclusion bodies from FD patients with the IVS4 mutation exhibited significant accumulation of inflammatory markers and oxidative stress markers. This observation suggests that early-stage Gb3 accumulation can induce significant cellular stress even before the manifestation of detectable pathological changes using current routine examinations, including HE staining and electron microscopic examination. The presence of inflammatory markers (NF-κB IL-18, and phospho-p42/44 MAPK) and the oxidative stress marker (iNOS) indicates an activated inflammatory response and increased oxidative stress in FD patients. These findings highlight the potential mechanisms underlying cellular damage in FD and provide future possible targets for therapeutic interventions, such as antioxidant supplementation or immunomodulatory therapy, especially in the early stages of the disease before expensive ERT or chaperone therapy becomes available.

Furthermore, the myocardial biopsies from G3Stg/GLAko mice and three FD patients without typical FD pathological findings demonstrated not only significant cellular stress but also significant fibrosis, despite the absence of inclusion bodies. Fibrosis is a hallmark of cardiac remodeling and can lead to impaired cardiac function [9]. The presence of fibrosis in these samples suggests that irreversible damage can occur in FD patients before the appearance of typical Fabry pathological changes. This finding highlights the importance of early intervention and treatment to prevent irreversible damage and improve patient outcomes.

Regarding the formation of Gb3 inclusion bodies, we believe it to be a result of Gb3 crystallization, which requires a concentration of Gb3 beyond saturation. Therefore, it is plausible that the appearance of inclusion bodies represents a later stage in the accumulation of Gb3. Our study indicates that before the manifestation of inclusion bodies, the intracellular Gb3 level might already be sufficiently high to induce significant inflammatory stress and irreversible tissue damage. These findings have important implications for the understanding of FD progression and the interpretation of histopathological examinations. The routine focus on inclusion bodies (lamellar myelin bodies) alone may underestimate the extent of cellular stress and damage present in FD cardiomyocytes. By solely relying on the presence of inclusion bodies (lamellar myelin bodies) as a diagnostic criterion, the disease may go undetected or be diagnosed at a later stage when irreversible damage has already occurred.

Due to the high prevalence of cardiac-type Fabry disease in Taiwan and our extensive series of myocardial biopsies, we had the opportunity to obtain these three precious biopsy samples. These samples are characterized by exhibiting significant Gb3 accumulation, as observed through immunostaining, yet did not display the typical pathological changes associated with Fabry disease. Interestingly, our findings in these biopsy samples were remarkably similar to those observed in 37-week-old G3Stg/GLAko mice, which corresponds to the human age range of 30 to 40 years. This is the period when patients with IVS4 FD typically develop initial cardiac symptoms. In both the mice and our three IVS4 FD patients who did not exhibit typical pathological changes but showed detectable Gb3 accumulation through immunostaining, we observed significant cellular stress and irreversible damage in the myocardial biopsies. These consistent results not only confirm that early-stage Gb3 accumulation can impair myocardial cells and cause irreversible damage but also highlight the potential of G3Stg/GLAko mice as a valuable research model for studying the impact of early Gb3 accumulation on the heart, especially when this specific type of human myocardial biopsies (without inclusion bodies) is challenging to obtain.

The limitations of this study included the inability to obtain myocardial biopsies from healthy human controls for comparison with patients, and the insufficient quantity of myocardial biopsy samples obtained from IVS4 patients for conducting Western blotting analysis. To address these constraints, we employed fibroblasts and a symptomatic mouse model of Fabry disease to perform multiple immunofluorescence staining and Western blot analyses. Importantly, we demonstrated that the immunofluorescence staining results from the mouse cardiac tissue were consistent with those observed in the patient cardiac biopsies. The concordance between the mouse model and patient samples was evident across several key inflammatory markers. For instance, both exhibited significantly elevated levels of globotriaosylceramide (Gb3) compared to their respective controls (Figure 1a, b and Figure 2a, b). Similarly, increased expression of inducible nitric oxide synthase (iNOS), interleukin-18 (IL-18), nuclear factor kappa B (NF-κB), and phosphorylated p42/44 mitogen-activated protein kinase (phospho-p42/44 MAPK) was observed in both the mouse model and patient samples (Figure 2c-j). Furthermore, the presence of early-stage fibrosis, as indicated by alpha-smooth muscle actin (α-SMA) immunofluorescence, was detected in both the G3Stg/GLAko mice and IVS4 Fabry disease patients (Figure 4a, b). The robustness of these parallel findings across different models and techniques strengthens the validity of our results. Therefore, we posit that the molecular and cellular events observed in our experimental models are representative of the pathological processes occurring in Fabry disease patients. This multi-model approach, despite the aforementioned limitations, provides a comprehensive view of the early inflammatory responses in Fabry disease.

## Conclusions

Our study challenges the conventional understanding of FD progression by highlighting the presence of significant oxidative stress, inflammation, and irreversible fibrotic changes before the appearance of Gb3 inclusion bodies. We propose that inclusion bodies may represent a later stage sign of Gb3 accumulation, occurring after intracellular Gb3 levels have reached a critical threshold. These findings call for a reevaluation of diagnostic criteria and emphasize the importance of early intervention strategies to mitigate cellular stress, inflammation, and fibrosis, ultimately improving the prognosis for patients with FD. Further research is warranted to validate these findings and explore alternative diagnostic markers and therapeutic approaches for the early detection and management of FD.

## Nonstandard Abbreviations and Acronyms

BD: Fixation/Permeabilization solution and Perm/wash solution
CTCF: Corrected total cell fluorescence
DAPI: 4,6-diamidino-2-phenylindole
ERT: Enzyme replacement therapy
FBS: Fetal bovine serum
FD: Fabry disease
Gb3: Gobotriaosylceramide
H&E: Hematoxylin and eosin staining
IF: Immunofluorescent
IL-18: Interleukin-18
iNOS: Inducible nitric oxide synthase
LAMP-1: Lysosomal-associated membrane protein 1
LVH: Left ventricular hypertrophy
MAPK: mitogen-activated protein kinase
NF-κB: nuclear factor-κB
OD: optical density
PBS: Phosphate-buffered saline
TgG3S: Transgenic human Gb3 synthase
WT: Wild type
α-Gal A: α-galactosidase A
GLAko: α-Gal A knockout
α-SMA: Alpha-smooth muscle actin

## Acknowledgments

We acknowledge the participation of study patients and their families.

## Authors’ contributions

Y.R.C. and D.M.N. was the major contributor in developing the concept. C.T.Y., and C.Y.H. analyzed the data. C.L.L. performed the experiments assisted by P.S.C., Y.Y.L. and Y.T.C..

C.L.L. wrote the manuscript with contributions from H.Y.L., Y.R.C. and Y.F.C.. All authors read and approved the submitted version.

## Availability of data and materials

All data generated or analyzed during this study are included in this published article.

## Ethics approval and consent to participate

The study protocol was conducted according to the Declaration of Helsinki and approved by the institutional reviews boards of Taipei Veterans General Hospital (VGHUST105-G7-6-1 and VGHUST106-G7-3-1). We obtained written informed consent from all participants.

## Consent for publication

Not applicable.

